# Glycan-Mediated Mechanosensing Regulates Megakaryocyte-Biased Hematopoietic Stem Cell Subsets

**DOI:** 10.1101/2025.01.25.634886

**Authors:** Alejandro Roisman, Leonardo Rivadeneyra, Lindsey Conroy, Melissa M. Lee-Sundlov, Natalia Weich, Simon Glabere, Shikan Zheng, Katelyn E. Rosenbalm, Mark Zogg, George Steinhardt, Anthony J. Veltri, Joseph T. Lau, Tongjun Gu, Hartmut Weiler, Ramon C. Sun, Karin M. Hoffmeister

**Affiliations:** Versiti Blood Research Institute and Translational Glycomics Center, Milwaukee, WI, USA; Department of Neuroscience, University of Kentucky, Lexington, KY, USA; Roswell Park Comprehensive Cancer Center, Department of Molecular and Cellular Biology, Buffalo, NY, USA; Department of Biochemistry & Molecular Biology, College of Medicine, University of Florida, Gainesville, FL, USA; Center for Advanced Spatial Biomolecule Research, University of Florida, Gainesville, FL, USA; Departments of Biochemistry and Medicine, Medical College of Wisconsin, Milwaukee, WI, USA

**Keywords:** glycosylation, B4GALT1, hematopoietic stem cell fate, chromatin rearrangement, hematopoiesis

## Abstract

Definitive hematopoietic stem and progenitor cells (HSPCs) development relies on intrinsic and extrinsic programs to meet homeostatic and stress-related demands. The comprehensive mechanisms governing HSPC’s fate remain poorly understood. Our study identifies B4GALT1, a glycosyltransferase essential for N-glycosylation, as a key modulator of HSPC lineage decisions. We demonstrate that B4GALT1 deficiency disrupts glycosylation patterns within the bone marrow (BM) niche, resulting in oncogenic glycan signatures and altered expression of Mucin13 in HSPCs. Loss of B4GALT1 expands HSPC pools and promotes megakaryocyte priming in HSPCs through transcriptional and chromatin modifications, enhancing the Wnt-Mucin13 axis. Mucin13, an oncogene characterized by aberrant glycosylation, underscores the critical role of B4GALT1 in sustaining BM glycosylation and mechanosensing, thereby regulating HSPC fate through functional, transcriptional, and chromatin dynamics. These observations provide insights into the impact of glycan structures on HSPC function, lineage reprogramming, and malignant transformation.

## INTRODUCTION

Blood cells originate from hematopoietic stem and progenitor cells (HSPCs) via a progression of progenitors such as multipotent progenitors (MPPs), common myeloid progenitors (CMPs), and megakaryocyte-erythroid progenitors (MEPs). Significant progress has been made in characterizing hematopoietic stem cells (HSCs) and multipotent progenitors (MPP1-4) with an evolving understanding of the HSPC spectrum ^1^. Recent research shows alternative pathways for HSCs to differentiate into megakaryocytes, with MPP2 directly driving megakaryocyte priming ^1,2^. Self-renewing, megakaryocyte-biased HSCs play a role in both normal and stress-induced hematopoiesis ^1,2^.

Protein glycosylation is governed by glycosyltransferases that reside in the secretory pathway. In a non-template fashion, glycosyltransferases expertly regulate the diversity of glycan structures found on cells in a remarkably well-defined manner ^3^. Although glycans have been recognized in hematopoiesis ^4,5^, their role in the differentiation and proliferation of HSCs into mature blood cells has not been extensively explored. In the bone marrow (BM) niche, the glycan-dense extracellular matrix and cell surfaces undergo alterations in response to chronic inflammation ^6^, aging ^7^, and hematological malignancies, including myelodysplastic syndrome (MDS) and myeloproliferative neoplasms (MPNs) ^4^. Thus, the interactome of cell-cell and cell-extracellular matrix (ECM) glycans likely regulates spatiotemporal dynamics to fine-tune HSPC fate decisions ^8^. Our findings underscore the critical role of the β-1,4-Galactosyltransferase 1 B4GALT1 in shaping the diverse glycosylation landscape of the BM niche and reveal the B4GALT1-Mucin13-Wnt-β-catenin axis as a crucial regulator of the megakaryocyte-primed stem cell pool multipotent progenitor (MPP) 2 population and LT-HSC expansion.

## RESULTS

### N-glycan diversity modulates marrow microenvironments and HSC function

HSPC function associates with intricate transcriptional regulators and specific stromal niches in the BM, dictating essential microenvironmental signals. Expanding HSCs primarily inhabit bone-remodeling cavities in the distal femur, reflecting a nuanced microenvironmental heterogeneity, which includes extracellular matrix members and glycans ^9^. Glycosylation provides eukaryotes with an elaborate and complex combinatorial system, creating a wide range of N- and O-glycan structures without genomic changes ^10^. B4GALT1 adds galactose (Gal) to N-acetylglucosamine (GlcNAc) residues to synthesize lactosamine (LacNAc) (**Figure 1A**), a required precursor for further modifications, including Lewis x (Le^x^/CD15) additions ^11^ crucial for cell-matrix interactions, homing, and myeloid lineage differentiation ^12–14^. B4GALT1 overexpression is linked to enhanced thrombopoiesis and thrombocytosis in MPN ^15^. In contrast, its deficiency leads to dysplastic megakaryocytes, impaired thrombopoiesis, and expanded HSPCs ^13^. Although the association between defective thrombopoiesis and aberrant β1-integrin function in the absence of B4GALT1 is established ^13^, the detailed mechanisms by which B4GALT1 influences HSPC regulation remain unclear.

**Figure 1:**
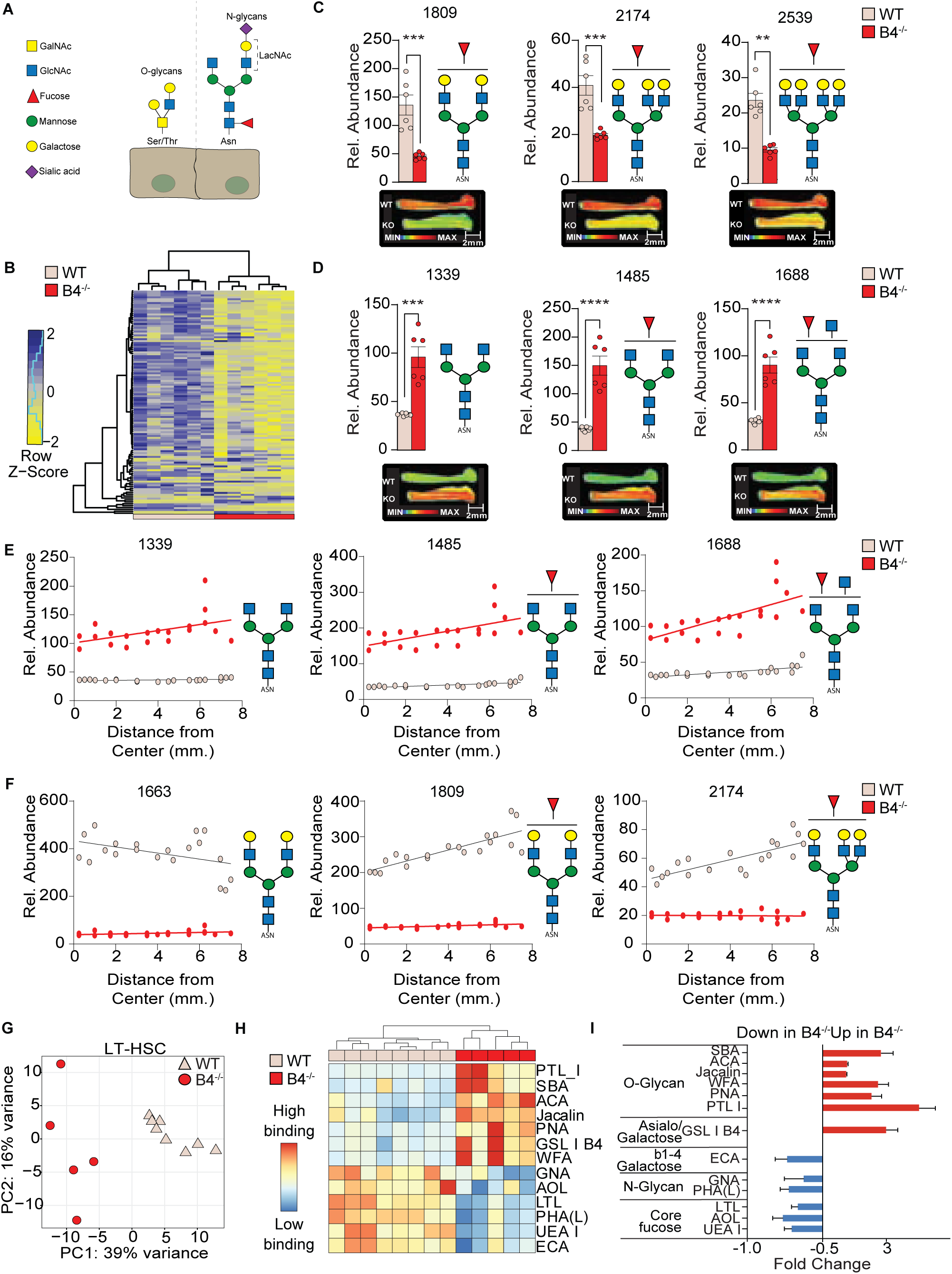
B4GALT1-dependent N-glycan diversity modulates marrow microenvironments and HSC function. **(A)** Schematic depiction of O- and N-linked glycan structures. N-linked glycans (N-glycans) are bound to proteins at asparagine (Asn) residues by an N-glycosidic bond (right). O-glycans are bound through O-glycosidic bond to serine (Ser) or threonine (Thr) (left). **(B)** Unsupervised clustering heatmap analysis of N-glycan structures in bone marrow compartments from control and B4^-/-^ femurs. Relative abundance of **(C)** galactosylated and **(D)** agalactosylated N-glycans in control and B4^-/-^ bone marrows. Representative N-glycan structures are depicted based on predicted monosaccharide composition from *m/z*. Spatial N-Glycan distribution of **(E)** galactosylated and **(F)** agalactosylated N-glycans measured from the center of the femur shaft to the distal compartments of control and B4^-/-^ specimens. **(G)** Principal Component Analysis (PCA) of 45 lectin binding data revealing distinct clusters for control and B4^-/-^ LT-HSCs. **(H)** Heatmap of differential lectin binding to control and B4^-/-^ LT-HSCs. **(I)** Relative lectin binding and their specificity to control and B4^-/-^ LT-HSCs is shown. Values represent individual samples ± SD, analyzed by Student t-test with Welch correction. Significance gradients are indicated as \**P* <.05; ** *P* <.01; *** *P* <.001; **** *P* <.0001).

To investigate this, we utilized B4GALT1 knockout mice (B4^-/-^ mice). By using PNGase F to specifically cleave N-glycans and Matrix-Assisted Laser Desorption Ionization-Mass Spectrometry Imaging (MALDI-MSI), we mapped the spatial N-glycan composition and distribution within age-matched control and B4^-/-^ BMs. Profound differences in N-glycan signatures were observed between controls and B4^-/-^ (**Figure 1B, Figure S1A**). Control BMs exhibited significant heterogeneity in the complex N-glycan spectra composition, marked by a higher abundance of galactosylated and core-fucosylated glycan epitopes within the 1600-2000 and 2000-2700 *m/z* range compared to B4^-/-^ BMs. Increased relative abundance for bi-antennary N-glycans, bisecting, and tetra-antennary N-glycans was observed in controls (**Figure 1C and figs. S1B and S1C**). Conversely, B4^-/-^ BMs displayed reduced diversity in complex N-glycans, with a 2-3-fold decline in N-glycans capped by galactose and a concurrent increase in agalactosylated N-glycans (*m/z* 1339, 1485 and 1688) (**Figure 1C and 1D and Figure S1C and S1D**). B4^-/-^ BMs demonstrated increased immature N-glycan epitopes, specifically pauci-mannose, commonly linked to aging and cancer (**Figure S1D**) ^16,17^.

Next, we assessed the N-glycan spatial distribution by examining the N-glycome from central to distal femur regions within the BM. N-glycan epitopes lacking galactose (i.e., *m/z* 1339, 1485, 1688) were of low abundance throughout all control femur regions. The abundance of specific glycan structures, such as core fucosylated, galactosylated bi-antennary, and bisecting N-glycan structures (i.e., *m/z* 1809 and 2174, respectively), significantly increased towards the distal femur region, a region is associated with high BM turnover and HSC expansion ^18^. In contrast, the non-fucosylated bi-antennary N-glycan (*m/z* 1663) demonstrated a marked decline in abundance towards the distal femur (**Figure 1E and 1F**). In B4^-/-^ BMs, MALDI-MSI showed a low abundance of core-fucosylated and galactosylated bi-antennerary and bisecting complex N-glycans across all examined regions, especially in the distal femur areas (**Figure 1F and Figure S1C**). Conversely, N-glycan epitope abundance lacking galactose (i.e., *m/z* 1339, 1485, and 1688) increased towards the distal femur region (**Figure1E and Figure S1D**).

While MALDI-MSI effectively mapped N-glycan distribution in the BM matrix, it lacks the resolution to discern specific N-glycan epitopes at a cellular level. Thus, we used a 45-member lectin array to probe glycan moieties of sorted control and B4^-/-^ LT-HSCs, and clear distinctions emerged between control and B4^-/-^ (**Figure 1G**). B4^-/-^ LT-HSCs demonstrated enhanced binding to lectins favoring O-glycans (PTL I, SBA, ACA, Jaclin, PNA, and WFA), and reduced affinity for N-glycan-associated lectins (GNA, AOL, LTL, PHA(L), UEA I, ECA). Most O-glycan-affinitive lectins specifically recognized cancer-associated motifs T-antigen (Galβ1→3GalNAcα1→Ser/Thr) (Jacalin, PNA, ACA) and Tn-antigen (GalNAcα1→Ser/Thr) (SBA, WFA) (**Figure 1H and 1I**). In contrast, B4GALT1-dependent structures, including lactosamine (ECA) and core-fucosylated glycans (AOL, LTL, UEA-I), were notably diminished ^3^ **(Figure1H and 1I**). These data show a decline in galactosylated complex N-glycans in B4^-/-^ samples, leading to a shift towards O-glycans linked to cancer pathways in the BM environment and on LT-HSCs ^3^, indicating that proper glycan distribution in the HSC compartment is maintained by B4GALT1-dependent N-glycosylation.

### Loss of N-glycans promotes HSPC proliferation without inflammation

We then thought to evaluate the effects of B4GALT1 deficiency on steady-state hematopoiesis. Consistent with prior findings ^13^, immunophenotypic evaluation showed a significant expansion of LT-HSCs (p=0.0061) (**Figure 2A**). Notably, we measured an increase in the megakaryocyte-biased LT-HSC population marked by CD41 expression (CD41^+^ HSCs) (**Figure 2B**) (p=0.0171), a LT-HSC subset that increases with aging or inflammation ^1,2^. We measured cytokine levels from BM supernatants to exclude inflammatory cues as triggers for the observed megakaryocyte-biased HSPC phenotype. Interleukins, chemokines, G-CSF, and GM-CSF levels in cell-free B4^-/-^ BM were comparable to or lower than controls (**Figure S2A-C**), denoting that B4GALT1 deficiency inherently drives proliferation of CD41-marked megakaryocyte-biased HSPCs and myeloid lineage differentiation, independent of inflammatory signals in the BM environment.

**Figure 2:**
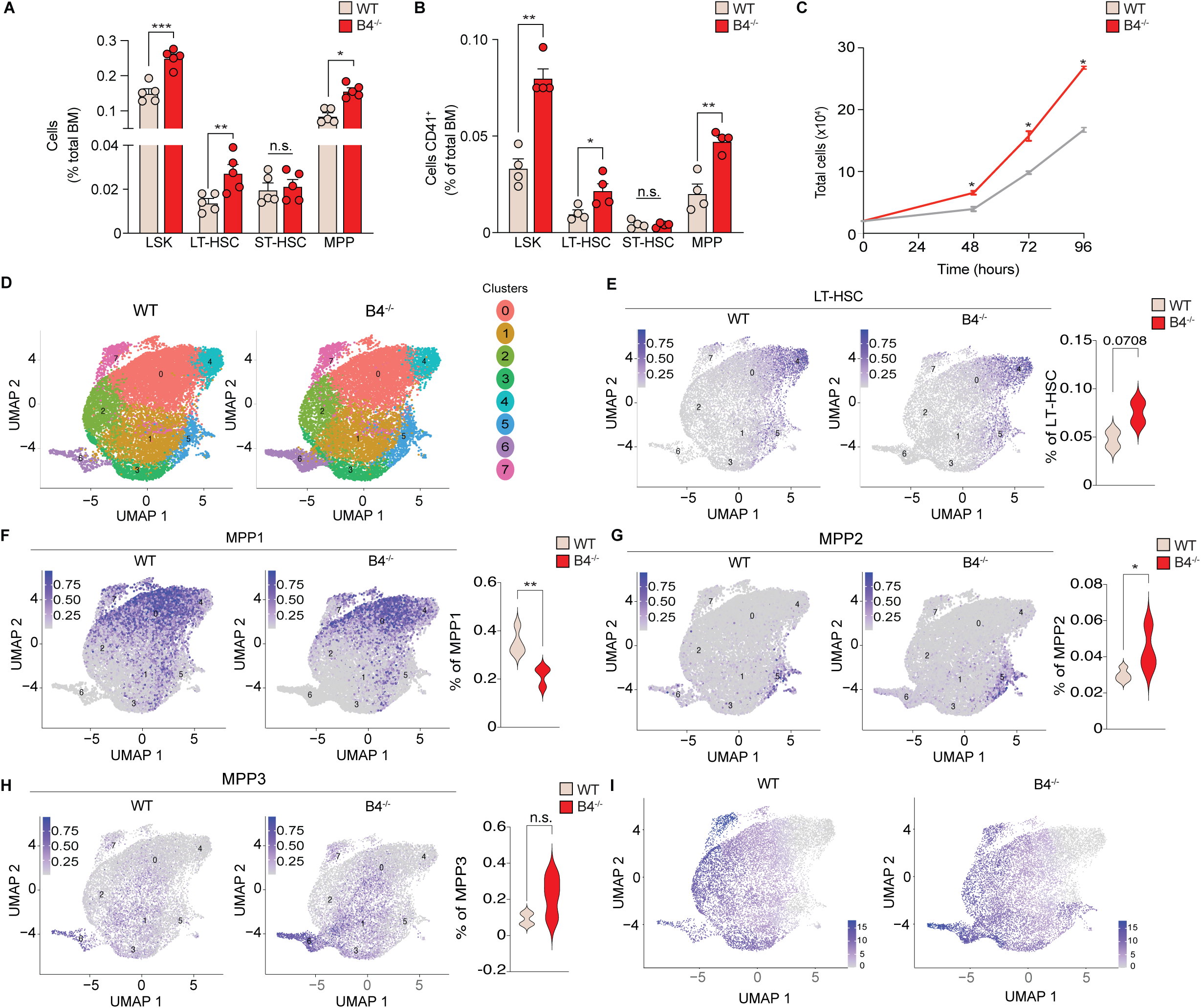
*B4galt1* loss induces hyper-proliferation and hematopoietic stem and progenitor (HSPC) expansion. **(A)** Immunophenotypic compositional flow cytometry analysis exhibiting HSPC expansion in B4^-/-^ bone marrows compared to control. **(B)** Flow cytometry immunophenotypic compositional analysis reveals the expansion of HSPC^CD41+^ in B4^-/-^ bone marrows and control. **C)** *In vitro* proliferation assay from sorted Lin^−^Sca-1^+^c-Kit^-^ (LSKs) cells from control and B4^-/-^ specimens **(D)** *In silico* identification of different transcriptional populations within all combined HSC and MPP subsets. **(E)** Uniform Manifold Approximation and Projection (UMAP) of transcriptionally defined LT-HSC compartments in B4^-/-^ specimens compared to control. **(F)** UMAP projections of transcriptionally defined MPP1 compartments in B4^-/-^ specimens compared to control. UMAP projections in transcriptionally defined MPP2 **(G)** and MPP3 **(H)** populations in control and B4^-/-^specimens. **(I)** Pseudotime potential in control and B4^-/-^ samples. Data are expressed as mean ± SD. Groups were compared using an unpaired Student’s t-test. Significance gradients are indicated as \**P* <.05; ** *P* <.01; *** *P* <.001; **** *P* <.0001. Color scales in **D-G** indicate expression levels of the gene signature utilized for the transfer learning approach. Color scales in **I** indicate trajectory in pseudotime projection.

*In-vitro* proliferation assays from sorted Lin^−^Sca-1^+^c-Kit^-^ (LSK) cells (**Figure S2D**), showed enhanced proliferation of B4^-/-^ LSK cells compared to controls (p<0.05, **Figure 2C**), with increased Lin^+^ (n = 4, p<0.05) (**Figure S2E**), and a tendency to increase CD41^+^, Ter119^+^ and CD11b^+^ cells (n = 2, **Figure S2F-H**). CD3^+^ and CD220 counts were indistinguishable from controls (n = 2, **Figure S2I**). This data reflects the HSPC bias toward megakaryocyte and myeloid differentiation observed in B4^-/-^ BMs ^13^ and supports the notion that B4GALT1 deficiency drives HSPC proliferation and myeloid differentiation, independent of inflammation or BM niche contributions.

### *B4galt1* absence expands transcriptionally defined LT-HSC and MPP2 pools

To explore whether global transcriptional profiles could elucidate the expansion observed in B4^-/-^ HSPCs, we performed single-cell RNA sequencing (scRNA-Seq) from sorted LSK cells (**Figure S3A**). Louvain clustering analysis of transcriptomes resulted in 8 different reproducibly distinct clusters (**Figure 2D and Table S1).** We then implemented a classification of single cells by transfer learning (CaSTLe) model to predict lineage classification based on existing transcript profiles ^19,20^ and identify hematopoietic lineages (LT-HSCs, MPP1-4) (**Figure S3B and Table S2**). Analysis of reported HSC signatures ^21,22^ further validated the lineage transcriptional identity (**Figure S3C and Table S3**), allowing us to recognize that LT-HSC pools were explicitly found in clusters 4 and 5 (**Figure S3D and Table S4**). Consistent with the immunophenotypic analysis, lineage transcriptional analysis identification ^19,20^ recognized LT-HSC expansion in B4^-/-^ cells (**Figure 2E and Table S5**) (p=0.07). Transposon tracing assays revealed that the MPP1 pool has a higher propensity to differentiate into lymphoid lineages ^20^. Analysis of the transcriptional potential across other HSPC compartments revealed a decrease in the MPP1 compartment in B4^-/-^ cells (**Figure 2F and Table S5).** The MPP2 compartment contains a high fraction of megakaryocyte-primed HSCs ^1,2,20^. Transcriptional output analysis uncovered a significant MPP2 expansion in B4^-/-^ samples (p<0.04) (**Figure 2G and Table S5**), consistent with megakaryocyte bias. No major changes in the MPP3 (**Figure 2H and Table S5)** or MPP4 (**Figure S3E and Table S5**) compartments were observed.

Hematopoiesis represents a continuum of differentiation with overlapping HSC subtypes ^23,24^ and early lineage priming in a subset of HSC ^20^. We inferred differentiation trajectories across hematopoietic lineages to delineate B4GALT1’s role in transcriptional priming during hematopoietic commitment ^25^. Notably, B4^-/-^ LT-HSCs from cluster 5 exhibited an increased pseudotime potential compared to controls (**Figure 2I**). Trajectory extremities indicated more efficient differentiation in B4^-/-^ LT-HSCs and MPP2 pools (**Figure S3F**).

### *B4galt1* loss enhances the megakaryocyte-primed LT-HSC population

Given the transcriptional and immunophenotypic megakaryocyte-priming, we next explored if *B4galt1* loss further expands additional megakaryocyte markers. *B4galt1* deficiency significantly augmented CD41 (*Itga2b*) expression in MPP2 (p<0.03) populations (**Figure S4A**) alongside an upregulation of other megakaryocyte-associated transcripts (*Pf4*, *GP9*, *Vwf*, *GP1ba*), indicating a shift towards a megakaryocyte-biased LT-HSC population (**Figure 3A**). Consistent with their lymphoid priming potential, the MPP1 population had decreased *Itga2b* expression, while there were no changes in MPP3 and 4 compartments (**Figure S4B**). LT-HSCs were transcriptionally defined to Clusters 4 and 5, overlapping with MPP2, which was confined exclusively to Cluster 5. This alignment likely indicates that this population corresponds to the Mk-bias HSCs. Transcriptomic analysis further revealed enrichment in genes associated with megakaryocyte development, particularly in cluster 5, indicative of megakaryocyte bias (**Figure 3B**), aligning with the immunophenotypically increased B4^-/-^ LT-HSC^CD41+^ population (**Figure 2B**). These data underscore B4GALT1’s selective role and targeted impact of B4GALT1 loss on directing LT-HSC and MPP2 progeny towards an expanded megakaryocyte-biased pool.

**Figure 3:**
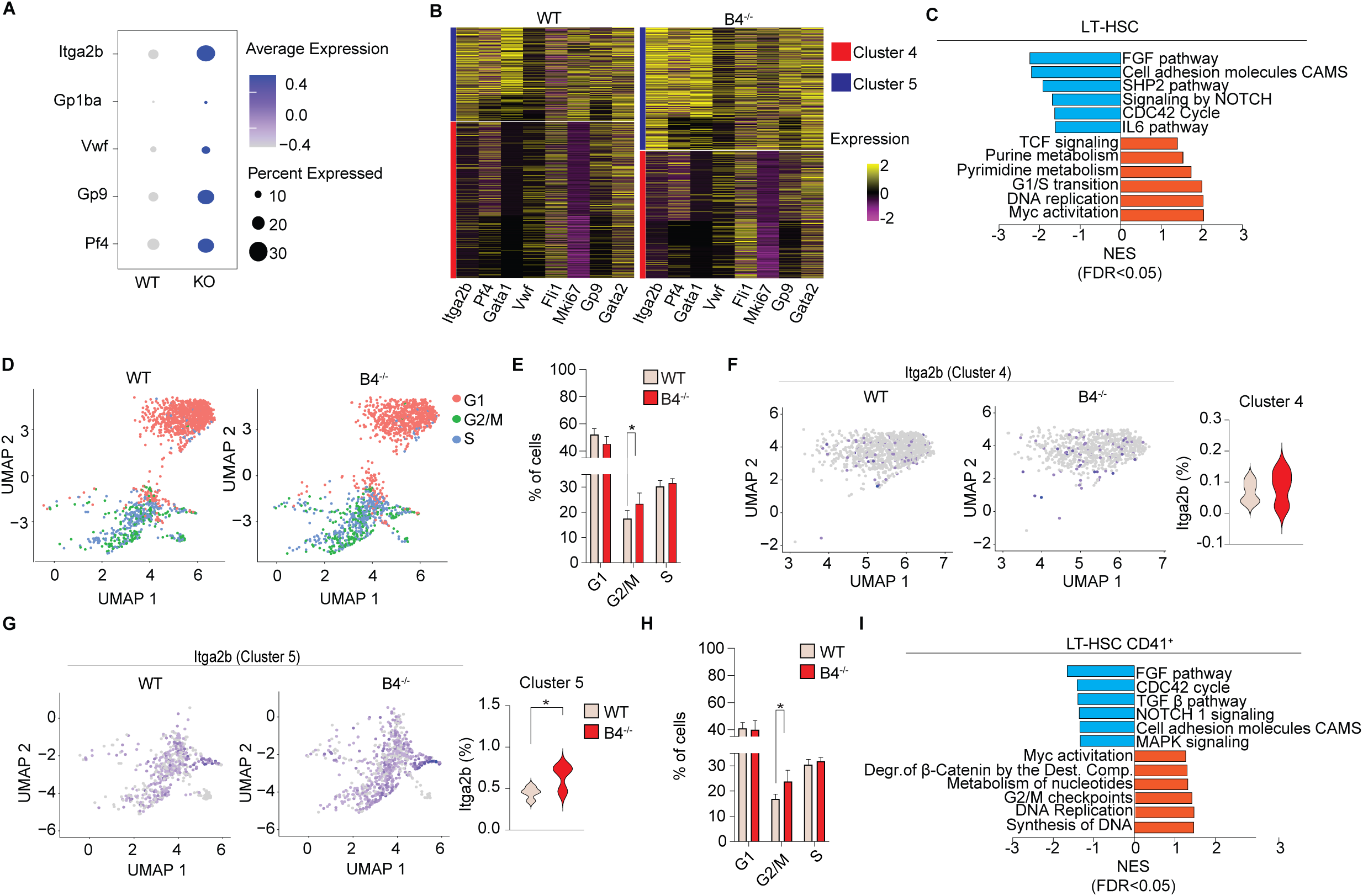
*B4galt1* loss alters cell cycle regulation and enhances the megakaryocyte-primed LT-HSC population. **(A)** Bubble plot representation of select megakaryocyte marker transcripts enriched in B4^-/-^ and control. **(B)** Heatmap representation of selected megakaryocyte-associated genes enriched in B4^-/-^ and control. **(C)** Bar plot of GSEA gene sets of select pathways that are significantly (FDR < 0.5) enriched in the LT-HSC compartment of B4^-/-^ and control specimens. UMAP distribution **(D)** and quantification **(E)** of HSPCs cell cycle states in B4^-/-^ and control. UMAP distribution and quantification of megakaryocyte marker *Itg2b* (CD41) in transcriptionally identified cluster 4 **(F)** and cluster 5 **(G)** LT-HSCs in B4^-/-^ and control samples. Color scales indicate *Itga2b* expression levels. (**H)** Cell cycle distribution of transcriptionally defined LT-HSC^CD41+^. **(I)** Bar plot of GSEA gene sets significantly (FDR < 0.5) enriched in the LT-HSC^CD41+^ compartment of B4^-/-^ and control samples. Data are expressed as mean ± SD. Groups were compared using an unpaired Student’s t-test. \**P* <.05.

### *B4galt1* deficiency enhances cell cycle activity in megakaryocyte-biased LT-HSCs

Gene Set Enrichment Analysis (GSEA) of B4^-/-^ LT-HSCs showed enrichment in metabolic and cell cycle regulatory pathways in B4^-/-^ LT-HSCs (**Figure 3C**) (FDR<0.05). In contrast, pathways crucial for HSC differentiation, such as cellular adhesion (CAMS) and NOTCH signaling pathways, were downregulated in B4^-/-^ LT-HSCs (FDR<0.05) (**Figure 3C and Tables S6 and S7**) ^26,27^. To decipher whether B4GALT1 influences HSC proliferation at the transcriptional level, we categorized LT-HSCs into the different cell cycle phases (**Figure 3D and Table S8**) ^28^. Cell cycle transcriptional analysis revealed an increase of G2/M phase in B4^-/-^ LT-HSC (p=0.0385), corresponding to clusters 4 and 5, and a slight decrease in cells in the G1 phase (**Figure 3E and Table S9).** We then explored whether changes in cell cycle activity of B4^-/-^ LT-HSCs correlated with variations in *Itga2b* (CD41) expression. The data shows no significant changes in cells from Cluster 4 (**Figure 3F**), but a significant increase in the fraction of cells expressing Itga2b within Cluster 5 (**Figure 3G**) (p=0.03), pointing to the exclusive enrichment of megakaryocyte-biased LT-HSCs in G2-M phase from cluster 5. Transcriptionally defined B4^-/-^ LT-HSC^CD41+^ cells also showed an increased presence in the G2/M phase (p<0.05) compared to controls (**Figure 3H**), which was accompanied by enrichment in cell cycle regulation pathways including DNA replication and G1 to S transition (FDR<0.05). Conversely, key pathways integral to HSC homeostasis and quiescence were depleted in the B4^-/-^ LT-HSC^CD41+^ cells, including NOTCH1 signaling, cell adhesion and TGF-β pathways, and MAPK signaling (FDR<0.05) (**Figure 3I and Table S10 and S11**), mimicking those changes observed in all LT-HSCs (**Figure 3C**). These results reveal that B4GALT1 deletion profoundly disrupts cell cycle dynamics and key cellular pathways in megakaryocyte-biased LT-HSCs, highlighting its essential role in maintaining HSC function and lineage fidelity.

### *B4galt1* loss induces Wnt-Myc signaling in megakaryocyte-primed HPSCs

We then sought to identify potential cause-effect relationships between transcriptional regulators and their targets ^29^. Causal network analysis revealed marked enrichment of canonical Wnt target genes in B4^-/-^LT-HSC cells (**Figure 4A and Table S12**). Myc, a crucial transcriptional regulator and downstream target of the Wnt/β-catenin signaling pathway, involved in HSC self-renewal and proliferation, ^27^ emerged as a top transcriptional regulator in B4^-/-^ LT-HSCs (**Figure 4B and Table S13)**. Transcriptional analysis further confirmed *Myc* among the top upregulated genes in B4^-/-^LT-HSCs (**Figure 4C** and **Table S7).**

**Figure 4:**
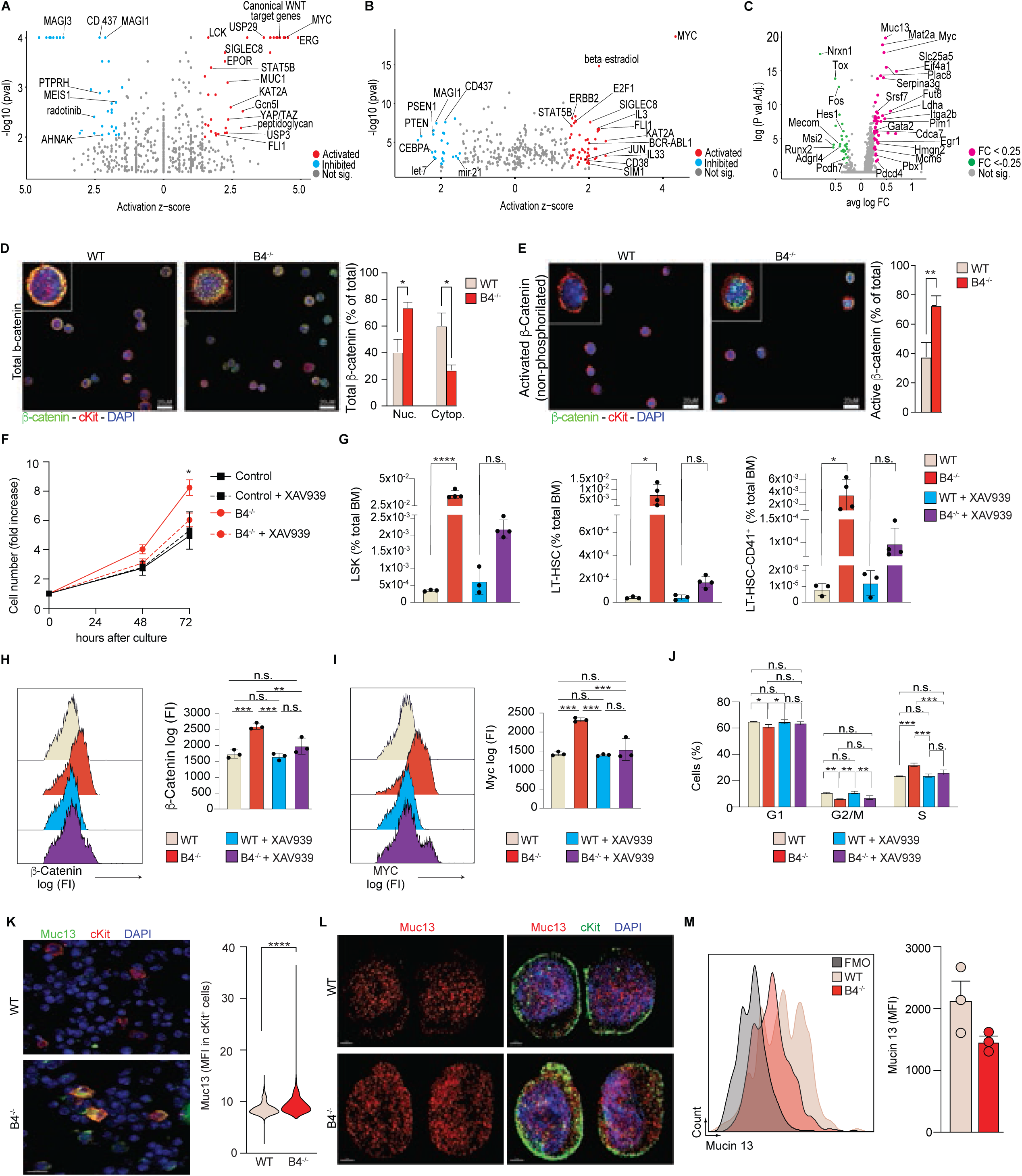
*B4galt1* loss induces increased Wnt-Myc signaling in megakaryocyte-primed HPSCs inducing expansion. **(A)** Volcano plot representation generated using Ingenuity Pathway Analysis software depicts differential activity between B4^-/-^ and control conditions, from the causal network analysis. **(B)** Volcano plot representation of Ingenuity Pathway Analysis reveals upstream regulator predictions in B4^-/-^ and controls. **(C)** Volcano plot representation of the log_2_ fold-change gene expression changes in B4^-/-^specimens compared to controls. **(D)** Immunofluorescence of sorted B4^-/-^ and control Lin^−^Sca-1^+^c-Kit^-^ (LSK) cells using an anti-β - catenin (total β-catenin, green) and anti-c-Kit (red) antibodies. Quantification of nuclear and cytoplasmic β-catenin localization is also shown (n=3). **(E)** Immunofluorescence of sorted B4^-/-^ and control LSK cells using anti-non-phosphorylated β-catenin (active β-catenin, green) and anti-c-Kit (red) mAbs. Quantification of nuclear β-catenin localization is also shown (n=3). Nuclei are shown using DAPI (blue). **(F)** Proliferation assay of cultured control and B4^-/-^ LSK cells in the presence of vehicle or the Wnt-pathway inhibitor XAV939 (n=3). **(G)** Phenotypic flow cytometry analysis using SLAM markers of sorted control and B4^-/-^ LSK cells treated with XAV939 or vehicle control. Quantification is also shown (n=3). **(H)** Representative flow cytometry histograms using an anti-β-catenin mAb (total β-catenin, left) of sorted control and B4^-/-^ LSK cells treated with XAV939 or vehicle control. Quantification is shown (right, n=3). **(I)** Representative flow cytometry histograms of Myc expression using an anti-Myc mAb (left) in sorted control and B4^-/-^ LSK cells treated with XAV939 or vehicle control. Quantification is also shown (right, n=3). **(J)** Cell cycle distribution quantification of sorted control and B4^-/-^ LSK upon treatment with XAV939 or vehicle control. **(K)** Representative immunofluorescence of Muc13 (green) and c-Kit (red) distribution in B4^-/-^ and control bone marrows (left). Quantification Muc13 and c-Kit colocalization (right). Muc13 mean fluorescence intensity (MFI) colocalized with c-Kit positive cells (cKit^pos^) is shown. **(L)** Representative immunofluorescence of sorted B4^-/-^ and control LSK cells using anti-Muc13 (red) and anti-c-Kit (green) antibodies. Nuclei are shown using DAPI (blue). **(M)** Representative flow cytometry histograms of Muc13 surface mean fluorescence (MFI) expression (left) in sorted B4^-/-^ and control LSK cells. FMO is shown for control. Quantification of Muc13 cell surface expression MFI in B4^-/-^ and control LSK cells (right) (n=3). All data are expressed as mean ± SEM. One-way ANOVA was used to compare each group. Significance is indicated as \**P* <.05; ** *P* <.01; *** *P* <.001.

Canonical Wnt signaling is regulated by β-catenin activity and depends on the inhibition of the destruction complex, allowing β-catenin cytoplasmic accumulation/nuclear translocation to initiate transcription of Wnt target genes ^30^. Immunoblotting of LSK cell lysates showed increased total β-catenin expression (**Figure S4C**), enhanced β-catenin (**Figure 4D**) and active, non-phosphorylated β-catenin (non-phosphorylated Ser33/37/Thr41) nuclear translocation in B4^-/-^ LSK cells (**Figure 4E**).

The Wnt-β-catenin pathway activates STAT3 and Wnt target genes, leading to metabolic reprogramming, Warburg effect, and increased lactate production ^31^. This further enhances Myc-driven glutaminolysis, essential for nucleotide biosynthesis in cell division. Higher Ldha expression in B4^-/-^ LT-HSC indicates metabolic shifts upon *B4glat1* loss (**Figure S4D and Table S6**). GSEA showed upregulation of pathways linked to cancer metabolism, nucleotide biosynthesis, and platelet activation, supporting Wnt-pathway activation in B4^-/-^ LT-HSCs (**Figure S4E and Table S10**). These results suggest that increased Wnt-Myc signaling in B4^-/-^ LT-HSC^CD41+^ cells promotes cell proliferation by modulating metabolic and nucleotide synthesis, predisposing these cells to megakaryocyte differentiation. To determine whether Wnt signaling enhances the proliferation of *B4galt1*-deficient cells, we conducted an *in-vitro* assay evaluating proliferation dynamics of control and B4^-/-^ LSKs with and without the Wnt inhibitor XAV939 ^32^. XAV939 treatment normalized B4^-/-^ proliferation to control levels (**Figure 4F**). Further analysis across different hematopoietic compartments showed that XAV939 treatment reduced the expansion observed in B4^-/-^ cells **(Figure 4G**). Additionally, XAV939 treatment decreased β-catenin and Myc levels in B4^-/-^ LSK cells (**Figure 4H-I**), aligning with findings that Wnt pathway inhibition reduces cell proliferation ^27^. Cell cycle analysis revealed that XAV939 treatment stabilized cell cycle distribution in B4^-/-^ cells to control levels (**Figure 4J**). These data strongly indicate that disrupted Wnt signaling is responsible for the excessive proliferation of *B4galt*-deficient hematopoietic cells, particularly in the LT-HSC^CD41+^ population.

### Mucin 13 is a potential Wnt/b-catenin signaling regulator in B4^-/-^ LT-HSC

Mucin 13 (Muc13), an oncogenic mucin ^33^, protects β-catenin from destruction complex mediated degradation by interacting with GSK-3β ^33^. Muc13, increasingly linked to cancer pathogenesis, affects cell growth, differentiation, and immune response ^34^. In cancer cells, Muc13 stabilizes β-catenin by inhibiting GSK-3β or binding directly to β-catenin ^35^, thus enhancing Wnt signaling and driving progression. In *B4galt1*-deficient LT-HSCs, possible upstream events regulating Wnt-β-catenin-Myc signaling include significant upregulation of *Muc13*, a transmembrane N-and O-glycosylated mucin (**Figure 4C, and Table S6 and S7**).

Bulk RNA-sequencing of sorted LT-HSCs confirmed a significant increase in Muc13 expression in B4^-/-^ LT-HSCs compared to controls (**Figure S4F Table S14**). BM sections showed increased co-localization of Muc13 with cKit^+^ cells in B4^-/-^ samples compared to controls (**Figure 4K**). In addition, immunofluorescence of isolated c-Kit^+^ cells revealed an increased Muc13 in the cytoplasm of B4^-/-^ LSK cells (**Figure 4L**), which was further validated by reduced Muc13 surface expression in B4^-/-^ LT-HSCs compared to controls (**Figure 4M**). Immunoblot analysis of sorted LSK cells displayed varied Muc13 glycoforms, with B4^-/-^ cells showing aberrant glycosylation patterns (**Figure S4G**), aligning with the altered N and O-glycosylation observed in B4^-/-^ (**Figure 1C-F and H).** This data suggests that *B4galt1* loss shifts Muc13 expression and glycosylation in LSK cells, including LT-HSCs, altering the Wnt-β-catenin-Myc signaling pathway.

### *B4galt1* deficiency drives megakaryocyte priming in HSPCs through transcriptional and chromatin changes

To determine if the functional impact of B4AGLT1 loss in HSCs is a cell-dependent effect, we generated a β4GALT1-Vav-cre mouse model carrying β4GalT1fl/fl LoxP sites on exon 2 to delete B4AGLT1 specifically from HSCs (B4^HSC-/-^). As described in B4^-/-^ mice ^13^, B4^HSC-/-^ mice have normal red blood and total white blood counts but display severe thrombocytopenia ^12,13^ (**Figure S5A**). The differential blood count showed an increase in neutrophil and monocyte numbers, while lymphocyte levels were reduced by 50%, indicating a myeloid skewing, as previously described ^12,13^ (**Figure S5A**). Further immunophenotypic analysis showed a marked increase in the LSK, LT-HSC, and MPP compartments, indicating enhanced stem and progenitor activity. Additionally, there was a notable rise in megakaryocyte-biased stem and progenitor cell fractions, suggesting a shift towards megakaryopoiesis, with no increase in bone marrow megakaryocyte numbers (not shown) (**Figure 5A and 5B**). To determine the functional impact of B4AGLT1 loss on HSPC function, we performed a combined single-cell RNA- and ATAC-seq (Multiome) on control and B4^HSC-/-^ specimens. Transcriptional annotation and lineage classification prediction confirmed expansion of the MPP2 compartment exclusively (**Figure 5C-E and Figure S5B**). GSEA revealed that B4^HSC-/-^ LT-HSC and MPP2s were enriched in megakaryocyte transcriptional regulators, such as RUNX1/3 ^36,37^, cell cycle regulation and WNT signaling (**Figure 5F and 5G**). B4^HSC-/-^ displayed upregulated megakaryocyte-associated transcripts indicating a shift towards a megakaryocyte-biased LT-HSC population (**Figure 5H, 5I and Figure S5C)**. Given the specific transcriptional changes observed in B4^HSC-/-^, we analyzed the remodeling of the chromatin accessibility landscape. Single-cell assay for transposase-accessible chromatin with high-throughput sequencing (scATAC-Seq) revealed that B4AGLT1 caused substantial changes in chromatin accessibility **(Figure 5J)**. Chromatin-accessible regions were predominantly linked to myeloid differentiation, while regions with reduced accessibility were associated with cell adhesion and ECM regulation (**Figure 5K)**, consistent with prior data obtained in B4^-/-^ mice. These findings suggest that the absence of B4GALT1-dependent glycosylation drives a shift in chromatin dynamics, disrupting ECM regulation and promoting myeloid-biased developmental programs. Transcription factor (TF) binding sites analysis revealed that while B4^HSC-/-^ LT-HSC were enriched in TFs essential for stem cell regulation (NFY and Stat3) ^38,39^, while MPP2s were associated with enhanced megakaryocyte-priming TFs (Runx1 and Fli1) ^37,40^ (**Figure 5L, Figure S5D)**. B4^HSC-/-^ MPP1, 3 and 4 subsets were enriched in TFs associated with immune differentiation, cell proliferation and T-cell regulation ^41–43^ (**Figure S5E).** Differentially accessible regions (DAR) in total B4^HSC-/-^ LSKs were associated with hematopoietic regulation, myeloid differentiation, and megakaryocyte developmental regulators, such as ERK1/2 ^44^ (**Figure 5M, 5N and Figure S5F**). Chromatin accessibility distribution from B4HSC^-/-^ revealed that LT-HSC which correlate with Cluster 0, MPP1s associated with Clusters 1-7, and MPP2s related to Cluster 5, represented the lineages exhibiting the highest number of gained peaks (**Figure S5G**), including an increase in chromatin accessibility at the Muc13 locus (**Figure S5H**). These findings show that B4GALT1 loss predominantly disrupts chromatin dynamics in LT-HSC and MPP2 compartments, reprogramming their commitment toward megakaryocyte development. Thus, cell-intrinsic B4GALT1-dependent glycosylation is essential for maintaining the immunophenotypic, transcriptional, and chromatin states required for balanced HSPC function. The loss of B4GALT1 reprograms HSPCs, driving accelerated commitment to the megakaryocyte lineage and skewing hematopoiesis toward megakaryocyte production at the apex of the HSPC hierarchy.

**Figure 5:**
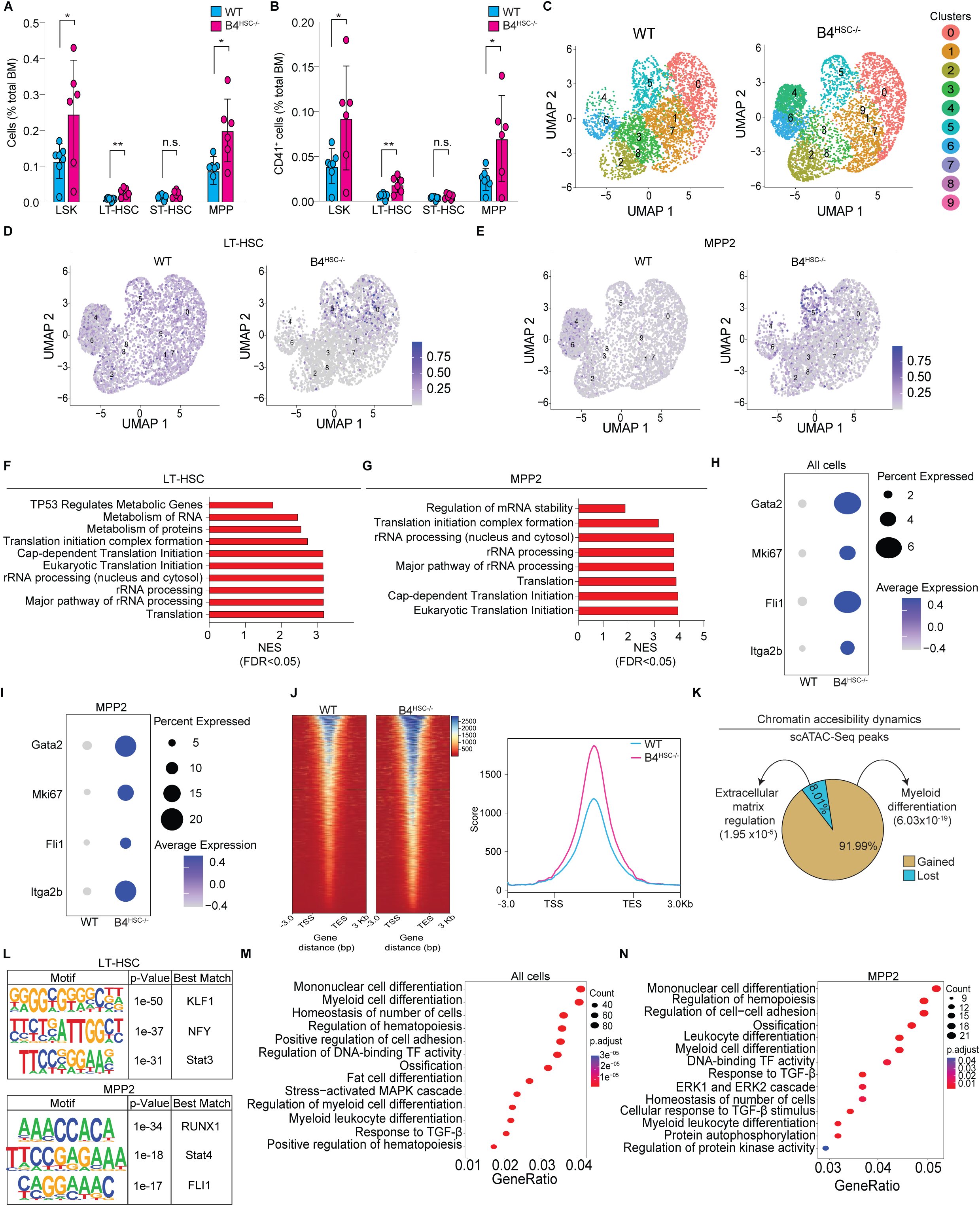
B4galt1 deficiency drives transcriptional and chromatin dynamics that favor megakaryocyte development in HSPCs. **(A)** Immunophenotypic compositional flow cytometry analysis of control (WT) and B4^HSC-/-^ bone marrows using SLAM markers. **(B)** Flow cytometry immunophenotypic compositional analysis reveals HSPC^CD41+^ expansion in B4^HSC-/-^ bone marrows compared to control (WT). **(C)** *In silico* identification of different transcriptional populations within all combined HSC and MPP subsets. UMAP projections of transcriptionally defined LT-HSC **(D)** and MPP2 **(E)** compartments in B4^HSC-/-^ specimens and controls. Bar plot of GSEA gene sets of select pathways that are significantly (FDR < 0.5) enriched in the LT-HSC **(F)** and MPP2 **(G)** compartments of B4^HSC-/-^ and controls. Bubble plot representation of select megakaryocyte marker transcripts enriched in B4^HSC-/-^ and controls **(H)** and MPP2 **(I)** B4^HSC-/-^ and controls. (**J)** scATAC-Seq signal heatmap representation in B4^HSC-/-^ and control (left) and density plot of scATAC-seq signal in B4^HSC-/-^ and control (right). (**K)** Percentage of gained and lost scATAC-seq peaks in B4^HSC-/-^ and control as well as gene ontology analysis and enriched transcription factor motifs of these peaks. (**L)** Motifs identified by hypergeometric optimization of motif enrichment (HOMER) of LT-HSC and MPP2 compartments of B4^-/-^ and control. Gene Ontology (GO) enrichment analysis of differentially accessible chromatin regions in B4^HSC-/-^ and control **(M)** and MPP2 **(N)** B4^HSC-/-^ and control specimens. Data are expressed as mean ± SD. Groups were compared using an unpaired Student’s t-test. Significance gradients are indicated as \**P* <.05; ** *P* <.01; *** *P* <.001; **** *P* <.0001.

## DISCUSSION

Our data highlights the critical role of B4GALT1 in regulating glycan structures within the BM niche, influencing LT-HSC and MPP2 commitment toward the megakaryocyte lineage. B4GALT1 deficiency disrupts fucosylated and sialylated N- and O-glycans in HSPCs, leading to oncogenic glycan patterns, including T and Tn antigens, and altering Mucin13 expression. Muc13, with its aberrant glycosylation, acts as an extracellular matrix sensor, triggering Wnt/β-catenin signaling hyperactivation. This cascade reprograms the epigenetic landscape of HSPCs, driving megakaryocyte-biased expansion.

Specific HSPCs compartments, particularly LT-HSC and MPP2, are critical drivers of megakaryocyte differentiation under steady-state conditions ^20^ with an increased propensity to develop into megakaryocytes following transplantation ^20^. Under stress conditions, the rapid increase in platelet counts, unlike other blood lineages ^45–47^, underscores the reliance on short-lived progenitors for emergency platelet production, highlighting the critical role of this lineage preference in rapid hematopoietic adaptation ^45,48^. Glycans are excellently suited to guide mechano-sensing to elicit rapid and emergency functional changes in protein structure and function in response to stress, given N-glycan rapid (hours) turnover on surface-expressed proteins ^49^. The remarkably similar phenotypic and genotypic effects of total and cell-specific B4GALT1 deletion highlight its indispensable role in these cell subsets. Recent studies identify galectin-1 as a key factor in myelofibrosis ^50,51^, binding lactosamine synthesized by B4GALT1 (**Figure 1A**). Our findings suggest that lactosamine-galectin-1 interactions are critical in regulating HSPC fate at a steady state. Further investigation is needed to clarify B4GALT1’s function in acute responses and under stress, including myelofibrosis.

Mucins, with their extended ectodomains, diverse domains, and variable glycosylation, are versatile glycoproteins evolved to protect exposed surfaces ^52–54^. Tumor cells exploit these altered mucin attributes to drive growth, proliferation, interaction with the extracellular matrix, and metastasis ^52–54^. Transmembrane mucins, including Muc1 and Muc13, have highly glycosylated extracellular domains that form protective mesh structures. In contrast, their intracellular domains regulate cell-cell interactions, proliferation, and apoptosis, functioning as external environment sensors ^55^. Muc13 is a key deregulated and aberrantly glycosylated mucin in B4^-/-^ HSPCs with increased intracellular levels, an oncogenic mucin ^35,56^. Muc13 protects β-catenin from destruction complex mediated degradation by interacting with GSK-3β ^33^. Muc13’s overexpression in leukemic models ^57^ and its association with increased Wnt signaling in the absence of B4GALT, suggests a direct role in promoting cell proliferation and malignant transformation ^58^. Regulation of canonical Wnt signaling entails disassembling the destruction complex within cells ^59^, activating β-catenin in HSCs, supporting their expansion ^60^, immature state preservation ^61^, and trilineage reconstitution ^27^. HSCs lacking B4GALT1 showed enhanced Wnt/β-catenin and Myc activity, and inhibiting Wnt signaling stabilized their expansion and proliferation, indicating a cell-autonomous increase in Wnt activity regulated by B4GALT1. Muc13 likely plays a regulatory role in HSPCs by acting as a sensor of the bone marrow environment, influencing key signaling pathways like Wnt/β-catenin to maintain the balance between different progenitor cell populations. Specifically, Muc13 could help regulate the function and maintenance of short-lived progenitors, such as megakaryocyte-biased HSPCs, ensuring proper platelet production and hematopoietic responses in stress conditions or “emergency cues”.

Cell fate determination is shaped by environmental cues, signaling pathways, transcriptional regulators, and epigenetic mechanisms. Our findings uncover a glycan-dependent regulatory network that integrates transcriptional, environmental, epigenetic, and mechanosensing controls to regulate HSPC function. B4GALT1 loss disrupts glycan metabolism and chromatin dynamics, increasing accessibility in regions linked to myeloid differentiation and reducing accessibility associated with cell adhesion and ECM regulation. These changes promote myeloid development and differentiation while driving transcriptional shifts that enhance megakaryocyte priming. Aberrant megakaryopoiesis, driven by defective differentiation of HSPCs rather than inherent malignancy in megakaryocytes, is a key factor in MPN development, including myelofibrosis ^62,63^. Considering *i)* Wnt/β-catenin aberrant signaling and associated risk of hematological malignancies ^64–67^, *ii)* the observed overexpression of B4GALT1 in AML models, ^68,69^ and *iii)* the transcriptional upregulation of B4GALT1, β-catenin, and Muc13 in MPN patient-derived HSPCs, especially those with JAK2^V617F^ mutations ^68,70^, it is plausible that aberrant glycosylation impairs Muc13 sensing ability and enhances the malignancy potential of megakaryocyte-biased HSPCs. The lack of targeted therapies against megakaryocytes ^71^ and megakaryocyte-biased HSCs highlights the need for new therapeutic strategies. CA19-9, the FDA-approved prognostic marker for pancreatic cancer, is a glycan antigen (sialyl Lewis^a^) on mucins like Muc1 ^72^. Muc1-based therapies are in preclinical and clinical trials ^73,74^. While CA19-9 expression by Muc13 is currently unknown, it is plausible that similar diagnostic tools and therapies could be developed to target and regulate hematopoietic cancer-associated aberrantly glycosylated HSPCs. B4GALT1 and Muc13 could be valuable targets for improving MPN treatment by modulating specific HSC and megakaryocyte functions.

In summary, our findings pioneer understanding the glycan-mechanosensing hierarchical role in the bone marrow, placing B4GALT1 as a key regulator of megakaryocyte-primed HSCs and glycan-niche diversity within hematopoiesis. We uncover the B4GALT1/Wnt/Muc13 axis as a mechanotransductive sensing pathway that governs the cellular and microenvironmental interface, regulating chromatin and transcriptional dynamics to promote HSC exit and megakaryocyte priming. Our integrated approach, combining MS-based glycomics, functional biology, and single-cell analysis, advances our understanding of HSC regulation and reveals clinically actionable pathways for disease treatment.

## Supporting information

Supplementary Materials and Other Supporting Files.

## RESOURCE AVAILABILITY

### Lead Contact

Requests for further information and resources should be directed to and will be fulfilled by the lead contact, Karin M. Hoffmeister.

### Materials availability

All materials used in this study are available upon reasonable request.

### Data and code availability

The datasets generated during this study are available in the Gene Expression Omnibus (GSE266314). This includes single-cell RNA-sequencing (scRNA-seq) datasets (GSE264078), bulk RNA sequencing (GSE263934), and single-cell Multiome (scMultiome) datasets (GSE283442).

## ACKNOWLEDGMENTS

We thank all Hoffmeister and Falet Lab members for their thoughtful discussions and suggestions. We would also like to thank the Versiti-Blood Research Institute Histology Core Lab and Versiti-Blood Research Institute Flow Cytometry Core Lab for their support and assistance in performing this study. We thank Drs. Hervé Falet, and Robert Burns for their critical review of the manuscript and data. A.R. is funded by the National Institute of Health K12 Translational Glyc*O*mics Program for Career Development in Glycoscience. We acknowledge Grace Kelly, Marge Kipp, and Michael Nemeth for their helpful discussions and help with the experimental design and procedures. This work was supported by National Institutes of Health grants R01 HL089224 (K.M.H.), P01 HL107146 (K.M.H.), K12 HL141954 (K.M.H.) and R01AG066653, R01CA266004 (R.C.S).

## AUTHOR CONTRIBUTIONS

Conceptualization: A.R., L.R., K.M.H. Methodology: A.R., L.R., L.C., S.G., M.L.-S., M.Z., N.W., G.S. S.Z., A.V., T.G. Investigation: A.R., L.R, L.C., N.W., M.Z., K.E.R., G.S., S.Z., A.V. J.T.L., H.W., T.G., R.C.S. K.M.H. Visualization: A.R., L.R., L.C., S.G., N.W., K.E.R., G.S., S.Z., T.G. Funding acquisition: R.C.S. K.M.H. Project administration: A.R., L.R, K.M.H. Supervision: H.W., T.G., R.C.S. K.M.H. Writing – original draft: A.R., K.M.H. Writing – review & editing: A.R., L.R., L.C., M.L.-S., N.W., S.G., G.S., S.Z., A.V., J.T.L., H.W., T.G., R.C.S. K.M.H.

## Methods

### Mice

β4GALT1^-/-^ mice were provided by the Consortium for Functional Glycomics (www.functionalglycomics.org). Wild-type littermates were used as controls. Mice were maintained as single strains on both C57BL/6J (JAX #000664) and 129S1**/**SvImJ (JAX #002448)backgrounds. For experiments, these mice were bred to produce mixed background 129S1**/**SvImJ/C57BLl/6J KO mice, using only the first generation after the crossing, and wild-type littermates were used as control. β4GALT1^flox^ mice were produced for the lab by Cyagen Biosciences, Inc. Vav1-icre (B6.Cg-*Commd10^Tg^*(Vav1–icre)*^A2Kio^*/J; JAX #008610) mice were acquired from Jackson Laboratories. Mice were maintained and treated as approved by the Institutional Animal Care and Use Committee of the Medical College of Wisconsin Committee according to National Institutes of Health standards as outlined in the Guide for the Care and Use of Laboratory Animals.

### Study approval

All experimental procedures involving animals complied with all relevant ethical regulations applied to using small rodents and with approval by the Animal Care and Use Committees (IACUC) at the Medical College of Wisconsin (Protocol No. AUA00005595).

### Bone marrow isolation, flow cytometry, and fluorescence-activated cell sorting (FACS)

Bone marrow (BM) cells were obtained by flushing mice tibia and femur shafts in 1×PBS supplemented with 3% FBS and 5 mM EDTA (flushing buffer) through a 70 m filter, followed by erythrocyte lysis 1× RBC Lysis Buffer (eBiosciences).

Cells were then stained in cold flushing buffer using the following antibodies: lineage cocktail (containing CD3e, CD5, Ter-119, Gr-1, Mac-1, and B220, eBioscience), c-Kit (clone 2B8, eBioscience), Sca-1 (Clone D7, eBioscience), CD150 (Clone TC15-12F12.2, Biolegend), CD48 (clone HM48-1, Biolegend). DAPI (Invitrogen) was used in all experiments for dead cell discrimination. Cell populations identified by flow cytometry were defined as: hematopoietic progenitors (LSK) – Lineage^NEG^, cKit^POS^, Sca-1^POS^; LT-HSC - Lineage^NEG^, cKit^POS^, Sca-1^POS^, CD48^NEG^, CD150^POS^. HSPC flow analysis was performed using LSR II and analyzed with BD Diva software. Cell sorting was performed in a BD FACSMelody using purity mode. Post-acquisition data analysis was performed with either BDFACS Diva or FlowJo software v10. For Myc and B-Catenin quantification, cells were stained with Myc Alexa Fluor 488 conjugated antibody (9E10, Santa Cruz Biotechnology Inc.) and beta-Catenin eFluor^TM^ 660 conjugated Antibody (15B8, Thermo Fisher) after ethanol fixation and permeabilization. Cell acquisition was performed in an LSRII (BD) instrument and analysis was performed using FlowJo Software (FlowJo, LLC).

#### Proteome profiler2

Bone Marrow supernatants were generated by crushing cleaned femurs and tibias in 400 μl of PBS in a sterile mortar. Samples were centrifuged (600 × g for 10 min), protein levels were quantified using Pierce™ BCA Protein Assay Kit, and 100 μg of total protein was used. According to the manufacturer’s instructions, cytokines were determined using Proteome Profiler Mouse XL Cytokine Array (R&D systems). Quantification was done using ImageQuant(TM) TL (GE Healthcare, Chicago, IL, USA) and the cytokine levels were expressed as arbitrary units.

### Lectin array

Lectin arrays were performed by sorting 1000 LT-HSC into NP40 lysis buffer. Cell lysates were stained with 10ug of Cy3 using GE Healthcare LS Cy3 Mono-Reactive Dye Kit (PA 23001, GE Healthcare Science). Cy3 excess was eliminated using Zeba™ Spin Desalting Columns, 7K MWCO, 0.5 mL (89882, ThermoFisher). After clean-up, samples were incubated overnight onto GlycoTechnica’s LecChip™ (GlycoTechnica). The mean intensity of fluorescence of each lectin was determined using GlycoStationToolsPro3.0 and SignalCapture3.0 (GlycoTechnica). Data obtained from different arrays was normalized using R studio by applying quantile normalization.

### Chemicals and Reagents

High-performance liquid chromatography-grade acetonitrile, ethanol, methanol, water, and trifluoroacetic acid (TFA) were purchased from Sigma-Aldrich. The α-cyano-4-hydroxycinnamic acid (CHCA) matrix was purchased from Cayman Chemical. Histological-grade xylenes were purchased from Spectrum Chemical. Citraconic anhydride for antigen retrieval was obtained from Thermo Fisher Scientific. Recombinant PNGaseF Prime was obtained from N-Zyme Scientifics (Doylestown, PA, USA).

### Formalin-fixed paraffin-embedded slide preparation for MALDI-MSI

Formalin fixed-paraffin embedded (FFPE) blocks were sectioned, mounted on positively charged glass slides, and processed similarly as described ^75,76^. Slides were heated at 60°C for 1 hr. After cooling, tissue sections were deparaffinized by washing twice in xylene (3 min each). Tissue sections were then rehydrated by washing slides twice in 100% ethanol (1 min each), once in 95% ethanol (1 min), once in 70% ethanol (1 min), and twice in water (3 min each). Following washes, slides were transferred to a Coplin jar containing citraconic anhydride buffer for antigen retrieval and the jar was placed in a vegeTable steamer for 25 min. Citraconic anhydride buffer was prepared by adding 25 µL citraconic anhydride in 50 mL water and adjusted to pH 3.0 with HCl. After antigen retrieval, slides were dried in a vacuum desiccator before enzymatic digestion.

### N-glycan MALDI-mass spectrometry imaging

An HTX spray station (HTX) was used to coat the slide with a 0.2 ml aqueous solution of PNGase F (20 mg total/ slide). The spray nozzle was heated to 45°C with a spray velocity of 900 m/min. Following enzyme application, slides were incubated at 37°C for 2 hr in a humidified chamber, and dried in a vacuum desiccator prior to matrix application [α-cyano-4-hydroxycinnamic acid matrix (0.021 g CHCA in 3 ml 50% acetonitrile/50% water and 12 µL 25%TFA) applied with HTX sprayer]. For the detection of N-glycans, a Waters SynaptG2-Si high-definition mass spectrometer equipped with traveling wave ion mobility was used. The laser was operating at 1000 Hz with an energy of 200 AU and spot size of 75 µm, mass range is set at 500 – 3000m/z. Ion mobility setting were done according to previously established parameters ^77,78^ with a trap entrance energy of 2V, trap bias of 85V, and DC/exist of 0V. Wave velocity settings were set to: trap 9.6 m/s, IMS 4.6m/s, transfer 17.4 m/s. Wave height settings were set to: trap 4V, IMS,42.7, transfer 4V, additional settings are variable wave ramp down of 1400 m/s. Images of N-glycans were generated using the waters HDI software.

### Bulk RNA-sequencing

Isolated cells (from 40-1000) were provided as cell pellets to generate a cDNA library for whole transcriptome analysis (RNAseq). Library preparation followed manufacturer’s recommendations in the SMART-Seq v4 Ultra Low Input RNA kit (Takara). Briefly, cells were lysed, and cDNA was prepared with the locked nucleic acid technology, template switching oligo, and primers that target the polyA tail of mRNA. Sample quality was verified by high-sensitivity DNA fragment analysis (Agilent, Bioanalyzer) to ensure that cDNA peaks greater than 800bp had been established and yielded greater than 4ng. Samples (150-300pg in 75uL of elution buffer) were sheared to 200-500bp using the Covaris E210 (175 peak power, 10% duty, 200 burst cycle, 5 min, frequency sweeping mode). Final preparation and amplification of libraries was completed with the SMARTer ThruPLEX DNA-seq Kit (Takara) utilizing dual 8bp indexes. Final libraries were checked by fragment analysis (Agilent, Bioanalyzer), quantified, and pooled by qPCR (Kapa Library Quantification Kit, Kapa Biosystems). Samples were sequenced over 2 lanes on the HiSeq2500, run in rapid mode with 2×150bp read lengths captured.

Illumina sequence adapters were trimmed from raw fastq files using CutAdapt. Trimmed reads were pseudoaligned using Salmon v.0.11.2 with reference GRCm38 [salmon reference: pmid 28263959]. These read counts were then imported into R (v3.4.3; R Core Team 2017) and DESeq2 v1.38.3 for further analysis. FPKM values from the fpkm module within DESeq2 were used for visualization and comparison.

### Single-cell RNA-Sequencing

Bone marrow LSK cells were sorted from 4 control and 3 B4^-/-^ mice. Cells were prepped by checking concentration and high-viability determined using a hemocytometer. 10,000 cells were loaded on a 10X Genomics Chromium Single Cell Controller to generate our single cell libraries. Single-cell capture and cDNA and library preparation were performed with a Single Cell 3′ v2 Reagent Kit (10X Genomics) according to the manufacturer’s instructions. Sequencing libraries were loaded on an Illumina NovaSeq 6000 instrument with an S1 flowcell and paired-end sequenced with the following read lengths: read 1, 26 cycles; read 2, 98 cycles, and a sample index of 8bp with a target coverage of 50,000 reads per cell.

### Multiome single cell RNA and ATAC sequencing

Single-cell RNA sequencing (scRNA-seq) and single-cell ATAC sequencing (scATAC-seq) libraries were prepared employing the 10X Genomics Single Cell Multiome assay kit (10X Genomics; 1000230, 1000283, 1000494, 1000215, 1000212) following the manufacturer’s guidelines. Briefly, LSK cells from ^B4HSC-/-^ and control specimens were sorted using antibody cocktails described above (Bone marrow isolation, flow cytometry, and fluorescence-activated cell sorting (FACS)), rinsed with 0.04% BSA PBS, and 10,000 cells underwent the manufacturer’s nuclei preparation procedure. Then, about 4,000 nuclei were placed into a 10x Chromium X instrument. The remainder of the library preparation steps were conducted in accordance with the manufacturer’s protocol. Libraries were evaluated using an Agilent 4150 TapeStation, quantified utilizing a KAPA Library Quantification Kit, and sequenced on an Illumina Novaseq platform.

### Single-cell RNA-seq analysis

Raw sequencing reads were demultiplexed and mapped to the latest mouse genome (mm10, https://cf.10xgenomics.com/supp/cell-exp/refdata-gex-mm10-2020-A.tar.gz) using 10x Genomics Cell Ranger v7.0.0 ^79^. This processing yielded a UMI count matrix for each cell and gene and were used as input for the Seurat suite of tools version 4.3.0 ^80^ using the R statistical package version 4.2.2 ^81^. We filtered cells with > 20% mitochondrial reads, and the cells having divergent counts (reads) relative to features (genes) were removed according to a quality control workflow provided by the Harvard Chan Bioinformatics Core ^82^.

### Clustering

The Seurat suite of tools was used to perform cluster analysis. First, raw read counts were normalized using sctransform (Variance Stabilizing Transformations for Single Cell UMI, version 0.4.0) with the percentage of mitochondrial reads regressed out. In addition, the effects of cell cycle heterogeneity calculated by CellCycleScoring were regressed out as described previously ^83^. The resulting normalized datasets were integrated across samples using standard Seurat functions. Briefly, SelectIntegrationFeatures PrepSCTIntegration, FindIntegrationAnchors and IntegrateData were used to identify the anchor genes with default settings, and integration across samples was carried out based on the anchor genes. Dimension reduction was performed on the integrated dataset through Principal Component Analysis, and FindClusters was used to generate the clusters. Finally, 8 clusters were yielded from an optimized resolution of 0.2631579 which was selected after comparing silhouette scores of a serial set of resolutions ^84^. Cells were prepared for downstream analysis of cluster marker genes using PrepSCTFindMarkers.

### Transfer Learning

For our LSK cell dataset, cell lineage was predicted by a transfer learning approach called CaSTLe ^19^. Briefly, classification models were trained using a dataset from a previous study with known lineage origin, then the trained model was used to estimate the classification probability of cells in our dataset. A cell was assigned to a lineage with the highest classification probability^20^.

### MolO and Basu enrichment score calculation

Previous studies have demonstrated that two gene sets, MolO ^22^ and Basu ^21^, are relatively enriched in LT-HSCs. In our study, we examined the enrichment of the two gene sets in all the clusters. Genes from MolO and Basu were used as input to the Seurat AddModuleScore function to calculate an enrichment score for each cluster, which was visualized in all clusters.

### Cell cycle prediction

Using the cc.genes function from the Seurat package, a list of human genes enriched in cell cycling were converted to mouse symbols by gprofiler “gorth” function. The cell cycle enriched genes were input to Serurat’s CellCycleScoring function to predict cell cycle status for each cell. Each cell was classified as G2M, S, or G1 phase.

### Ingenuity Pathway Analysis (IPA)

Differential gene sets were input to IPA’s expression analysis program that yielded upstream regulators and causal network predictions. A cutoff of log2FC at +/-0.25, and adjusted p-value at 0.05 were used in the IPA analysis.

### RNA velocity analysis

We used the standard approach from Velocyto for the RNA velocity analysis. Briefly, loom files were generated using Velocyto ^85^ command-line function. These files were then converted into Seurat objects. Subsequently, the nearest neighbor graph and reduction slot of single-cell Seurat object were incorporated into Velocyto Seurat objects. Velocity computation was performed using RunVelocity function with the following parameters: deltaT = 1, kCells = 25, fit.quantile = 0.02. Distinct colors were assigned to different clusters for visualization purposes. Velocity Figures were generated using the velocyto.R function, show.velocity.on.embedding.cor with the following parameters emb = Embeddings(object = object, reduction = “umap”), vel = Tool(object = object, slot = “RunVelocity”), n = 200, scale = “sqrt”, cell.colors = ac(x = cell.colors, alpha = 0.5), cex = 0.8, arrow.scale = 3, show.grid.flow = TRUE, min.grid.cell.mass = 0.5, grid.n = 40, arrow.lwd = 1, do.par = FALSE, cell.border.alpha = 0.1, xlim = c(-15, 15), ylim = c(-15,15)).

### Trajectory/pseudotime analysis

Seurat objects were converted to cell_data_set objects. Partitions were assigned to the CDS object. Subsequently, cluster identification and UMAP visualization of the single-cell Seurat object were integrated into CDS objects. Trajectory computations were conducted using the learn_graph function from Monocle3 with following parameters: use_partition=TRUE. For visualization, trajectory Figures were generated using the plot_cell function with following parameters: color_cells_by = ‘cluster’, label_groups_by_cluster = FALSE, label_branch_points = FALSE, label_roots = FALSE, label_leaves = FALSE, group_label_size = 5. Pseudotime was computed using the pseudotime function from Monocle3. Pseudotime Figures were then plotted using the FeaturePlot function with the parameter feature=’pseudotime’ from the Seurat package.

### scMultiome analysis

Raw scRNA-seq and scATAC-seq reads were aligned to mm10 using Cellranger arc v2.0.2. Seurat v5.1.0 ^86^ and Signac v1.13 ^87^ were used for downstream analysis. All mitochondrial genes were removed from the scRNA-seq dataset. scRNA-seq assay were normalized and scaled using SCTransform v0.4, while scATAC-seq assay underwent normalization and latent semantic indexing (LSI) for dimensionality reduction. Integrative analyses were performed on both assays. To integrate the scRNA-seq and scATAC-seq assays, the Seurat Weighted Nearest Neighbor (WNN)^80^ approach was applied. The FindMultiModalNeighbors function was used to calculate WNNs by combining information from both modalities, giving more weight to RNA or chromatin data as needed. Clusters were then identified using the WNN graph, and UMAP was used to visualize joint cellular states. Differentially expressed genes (DEGs) and differentially accessible chromatin regions between KO and WT groups were identified using Wilcoxon Rank Sum test in Seurat^88^. Gene Set Enrichment Analysis (GSEA) was performed using clusterProfiler v4.10.1^89^, where genes were ranked by fold-changes. KEGG^90^, Reactome^91^, and Gene Ontology^92^ databases were used in the analysis of GSEA. For over-representation analysis, significant differential peaks were selected, and Gene Ontology database was applied. To control the false positive rate, multiple testing correction was applied using the Benjamini-Hochberg method to adjust the p-values obtained from the differential analysis, GSEA, and over-representation test. We set a significance threshold of adjusted P value at 0.05 to control the false discovery rate at 5%^93^. Motif analysis was performed using findMotifsGenome.pl from Homer and FindMotifs function from Signac. DotPlot and FeaturePlot were used for RNA-seq visualization. Deeptools v3.3.1^94^ was used for ATAC-seq visualization. Trajectory analysis was applied using Monocle3 ^95^.

### Immunoblotting

Protein lysates were subjected to gel electrophoresis (SDS–PAGE) and transferred to polyvinylidene fluoride (PVDF) membrane (BioRad, Hercules, CA, USA). Rabbit anti-CTTNB1 (clone CAT-15, catalog number 712700, Invitrogen) antibody and rabbit anti-GAPDH 1 antibody (clone 14C10, catalog number 402200, Cell signaling Technology) were used for western blot. Immunoreactive bands were detected by horseradish peroxidase-labeled streptavidin (1:3000, Cell Signaling Technology) or horseradish peroxidase-labeled secondary antibodies (BioRad), using enhanced chemiluminescence reagent (Millipore). Pre-stained protein ladders (BioRad) were used to estimate the molecular weights. Band intensities from individual western blots were quantified by densitometric analysis using ImageJ software.

### Immunofluorescence of mouse bone sections

Femurs were harvested and fixed into periodate-lysine-paraformaldehyde fixative (0.01 M Sodium-M-Periodate, 0.075M L-Lysine, 1% PFA) overnight at 4°C. Femurs were rehydrated into 30% sucrose in phosphate buffer for 48 hours, embedded in OCT (TissueTek®, Sakura Finetek USA, Torrance, CA, USA), and snap frozen in isopentane/dry ice mixture. Whole longitudinal single-cell-thick (7 μm) femoral cryosections were obtained using a Leica Cryostat and the Cryojane tape transfer system (Leica Microsystems, Wetzlar, Germany). Slides were thawed, permeabilized with T-TBS (0.1%), and blocked with 5% bovine albumin serum. Primary antibodies for detecting CTTNB1 (clone CAT-15, catalog number 712700, Invitrogen) and Non-phospho (Active) β-Catenin (Ser33/37/Thr41) (clone D13A1, catalog number 8814, Cell signaling Technology) were incubated at room temperature for 1 hour. Slides were washed, incubated with Image-IT signal enhancer for 30 minutes, and finally incubated with Alexa Fluor conjugated secondary antibodies (1:1000, Invitrogen). Immunofluorescence images were acquired by an Nikon Ti2 inverted laser scanning microscope (Olympus, Deutschland GmbH, Hamburg, Germany). For quantification, surfaces were created around the staining using Imaris software (Bitplane, Switzerland).

For LSK staining 15,000-25,000 cells were sorted into flushing buffer using FACSMelody sorter. Cell suspensions were centrifuged at 500rpm for 3 minutes using Cyto-tek cytocentrifuge model 4325 onto positively charged slides. Cells were allowed to air dry on the slide for 30 min then fixed in methanol for 15 min. Slides were blocked with tris buffered saline (TBS) with 0.1% Tween (T-TBS) and 5% Bovine Serum Albumin (BSA) for 1 hour and incubated with primary antibodies in T-TBS with 1% BSA overnight at 4C. Secondary staining was done using corresponding Alexa fluor plus secondary antibodies. Slides were mounted using ProLong™ Glass Antifade Mountant and visualized using an Olympus FV1000MPE Laser Scanning Confocal Microscop.

### In-vitro proliferation assay

LSK cells were sorted into StemSpan SFEM (STEMCELL technologies), spun down, and resuspended in 200ul of SFEM medium supplemented with SCF (50 ng/ml) and TPO (10 ng/ml) (all from Peprotech). To assess WNT pathway function in B4^-/-^ LSK cells, cells were treated with the inhibitor XAV939 (Catalog: 3748, Tocris) or vehicle (DMSO) at 0 and 72 hs. Cell number was evaluated using a hemocytometer using trypan blue to determine viability.

### Cell cycle analysis

Cell cycle profiles were assessed by flow cytometry after ethanol fixation and permeabilization, cells were stained with propidium iodide and analyzed using FlowJo Software (FlowJo, LLC).

### Statistical analysis

All experiments were performed at least in triplicate and data are represented as mean ± standard error of the mean (SEM). Numeric data were analyzed using one-way ANOVA analysis of variance followed by Bonferroni adjustment for multiple comparisons. Two groups were compared by the two-tailed Student’s unpaired t-test. The significance of data was assessed using the GraphPad Prism 5 software. Differences were considered as significant when p < 0.05. Different levels of significance are indicated as *p < 0.05, **p < 0.01, ***p < 0.001.

## Supplementary Information

### Content

- **Figures S1** to **S5.**
- **Document S1**. Tables S1 to S14. Excel file containing data too large to fit in a PDF file.
  - **Supplementary Table S1:** Differentially expressed genes obtained by Louvain clustering analysis of transcriptomes. Related to Figure 2 and supplementary Figure 3.
  - **Supplementary Table S2:** Gene expression signatures of cells classified as LT-HSC, STHSC, MPP as per Cell Assignment by Transcriptome Learning and Expression (CaSTLe) classification. Related to Figure 2 and supplementary Figure 3.
  - **Supplementary Table S3:** Comparison of Basu scores across clusters
  - **Supplementary Table S4:** Comparison of Molo scores across clusters
  - **Supplementary Table S5:** Classification of stem and progenitor compartments in control vs. B4^-/-^ LT-HSC. Related to Figure 2.
  - **Supplementary Table S6:** Gene set enrichment analysis. Related to Figure 3. Reported are the significant (FDR<0.05) GSEA results for the Wald statistic ranked lists of control vs. B4-/- LT-HSC.
  - **Supplementary Table S7:** Differentially expressed genes of control vs. B4^-/-^ LT-HSC. Related to Figure 3. Reported are the differentially expressed (p-adj<0.05, fold-change >2) genes of control vs. B4^-/-^ LT-HSC.
  - **Supplementary Table S8:** Cell cycle transcriptional markers employed to classify cell cycle status of LT-HSC from control vs. B4^-/-^ LT-HSC. Related to Figure 3.
  - **Supplementary Table S9:** Classification of control and B4^-/-^ LT-HSC according to their cell cycle distribution. Related to Figure 3.
  - **Supplementary Table S10:** Gene set enrichment analysis. Related to Figure 3. Reported are the significant (FDR<0.05) GSEA results for the Wald statistic ranked lists of control vs. B4-/- LT-HSC CD41+.
  - **Supplementary Table S11:** Differentially expressed genes of control vs. B4-/- LT-HSC. Related to Figure 3. Reported are the differentially expressed (p-adj<0.05, fold-change >2) genes of control vs. B4-/- LT-HSC CD41+.
  - **Supplementary Table S12:** Causal Network Analysis of upstream regulators in control vs. B4-/- LT-HSC. Related to Figure 4. Reported are the significant results obtained according to the differentially expressed genes in control vs. B4-/- LT-HSC.
  - **Supplementary Table S13:** Upstream Regulator Analysis determining likely regulators connected to differentially expressed genes observed from control vs. B4-/- LT-HSC. Related to Figure 4. Reported are the significant results obtained according to the differentially expressed genes in control vs. B4-/- LT-HSC.
  - **Supplementary Table S14:** Gene expression signatures of sorted LT-HSC in control vs. B4-/- LT-HSC. Related to Supplementary Figure 5.

