## Supplementary figures and images for "Glycan-Mediated Mechanosensing Regulates Megakaryocyte-Biased Hematopoietic Stem Cell Subsets"

### Figure S1

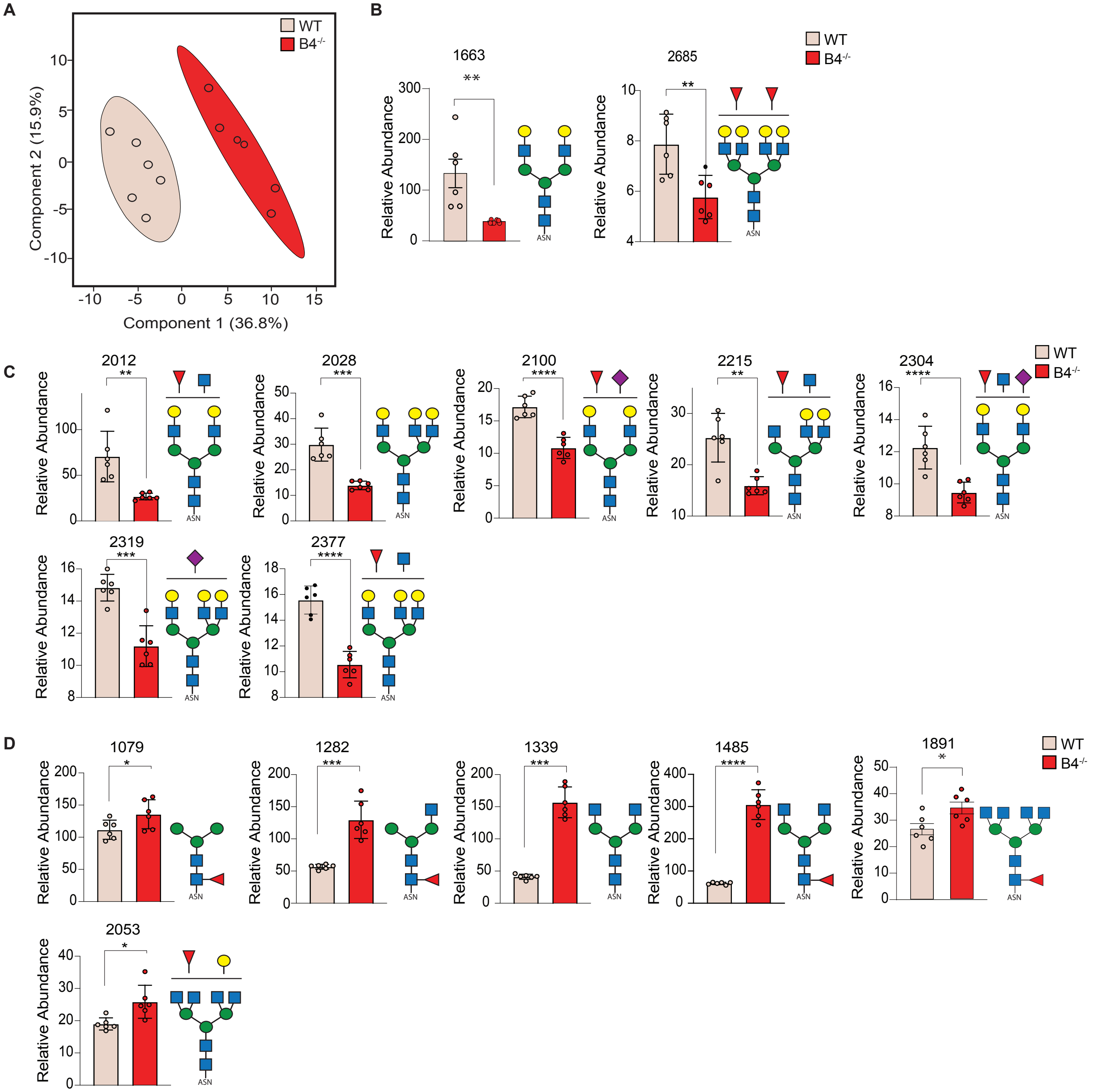

### Figure S2

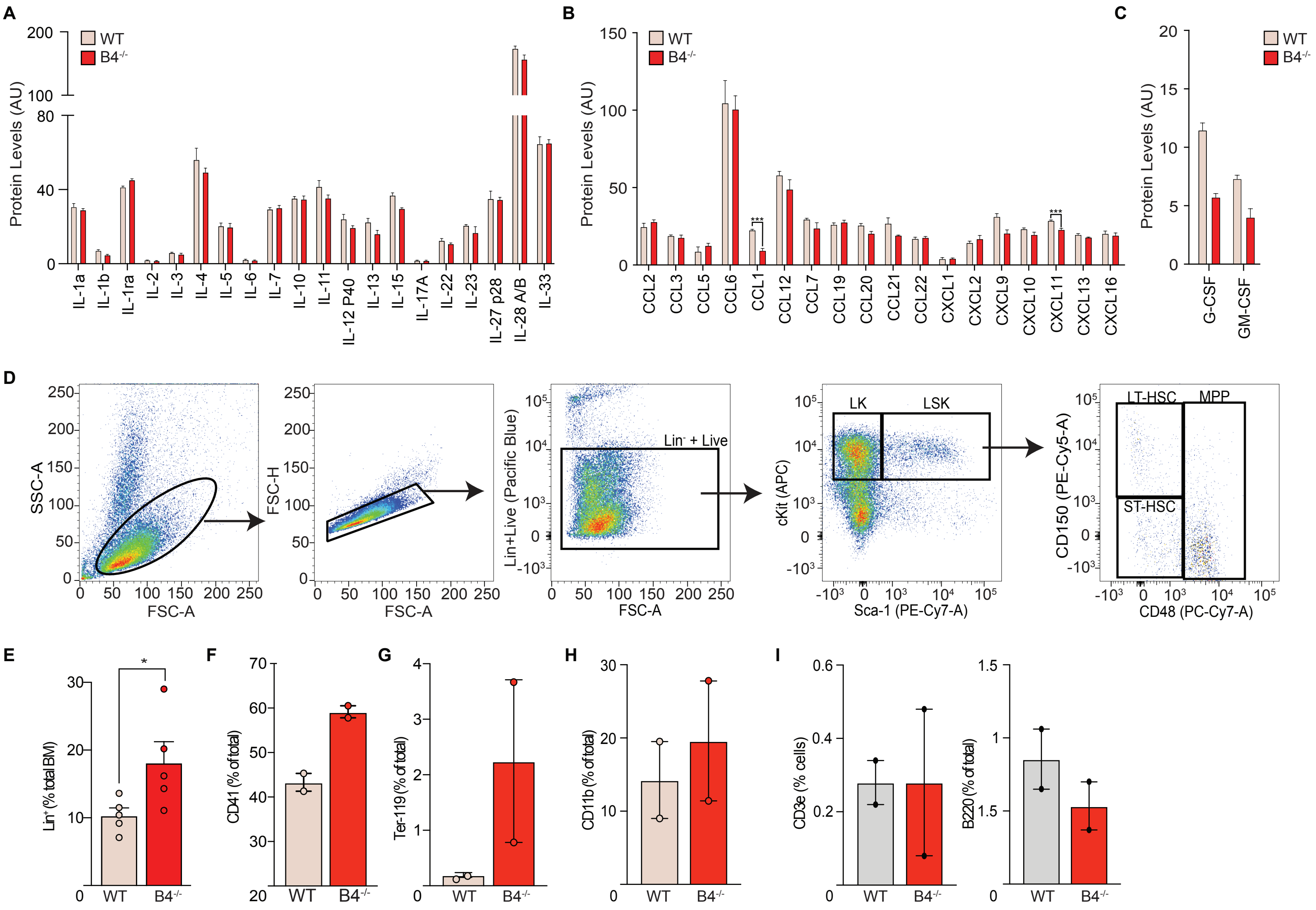

### Figure S3

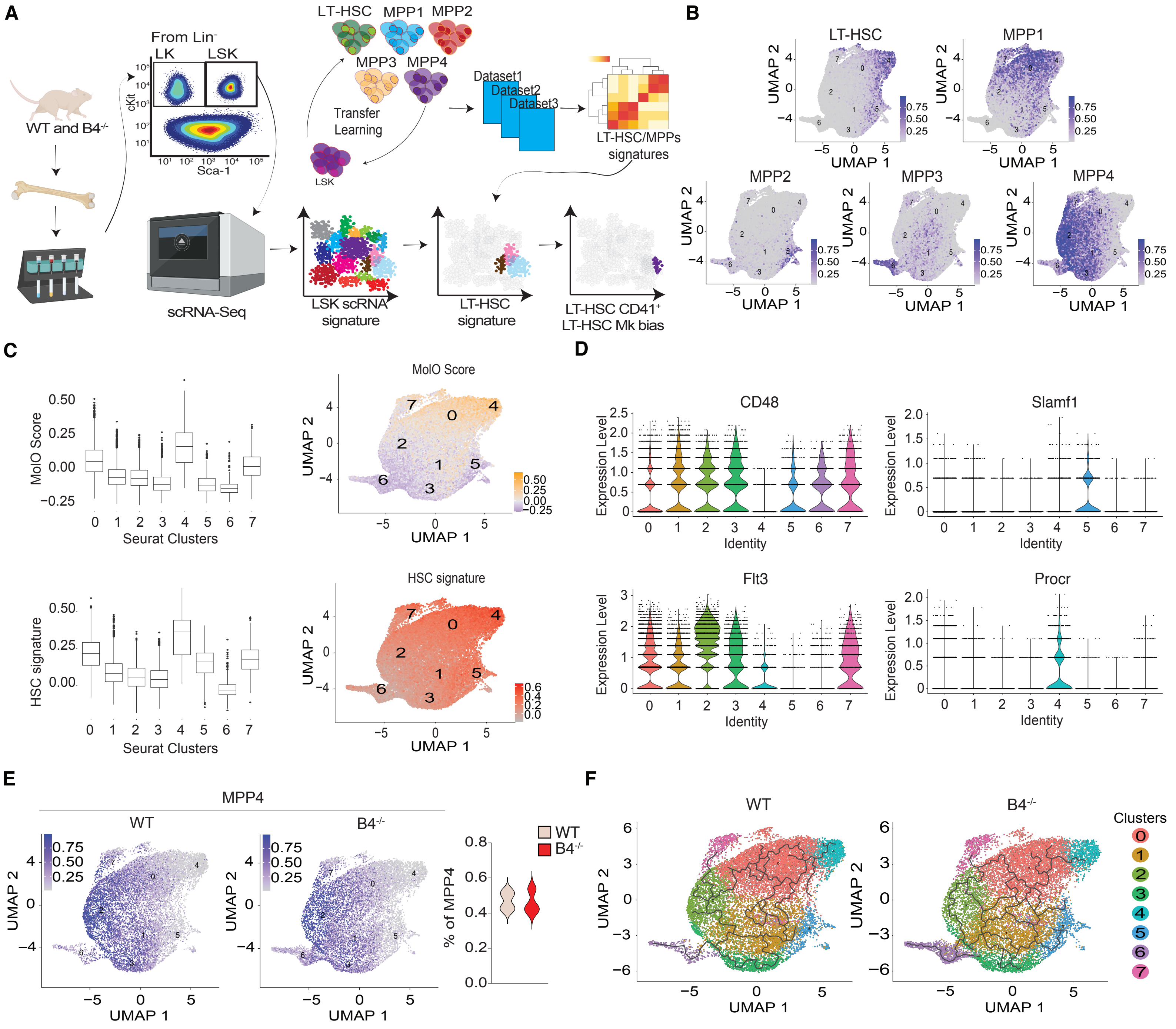

### Figure S4

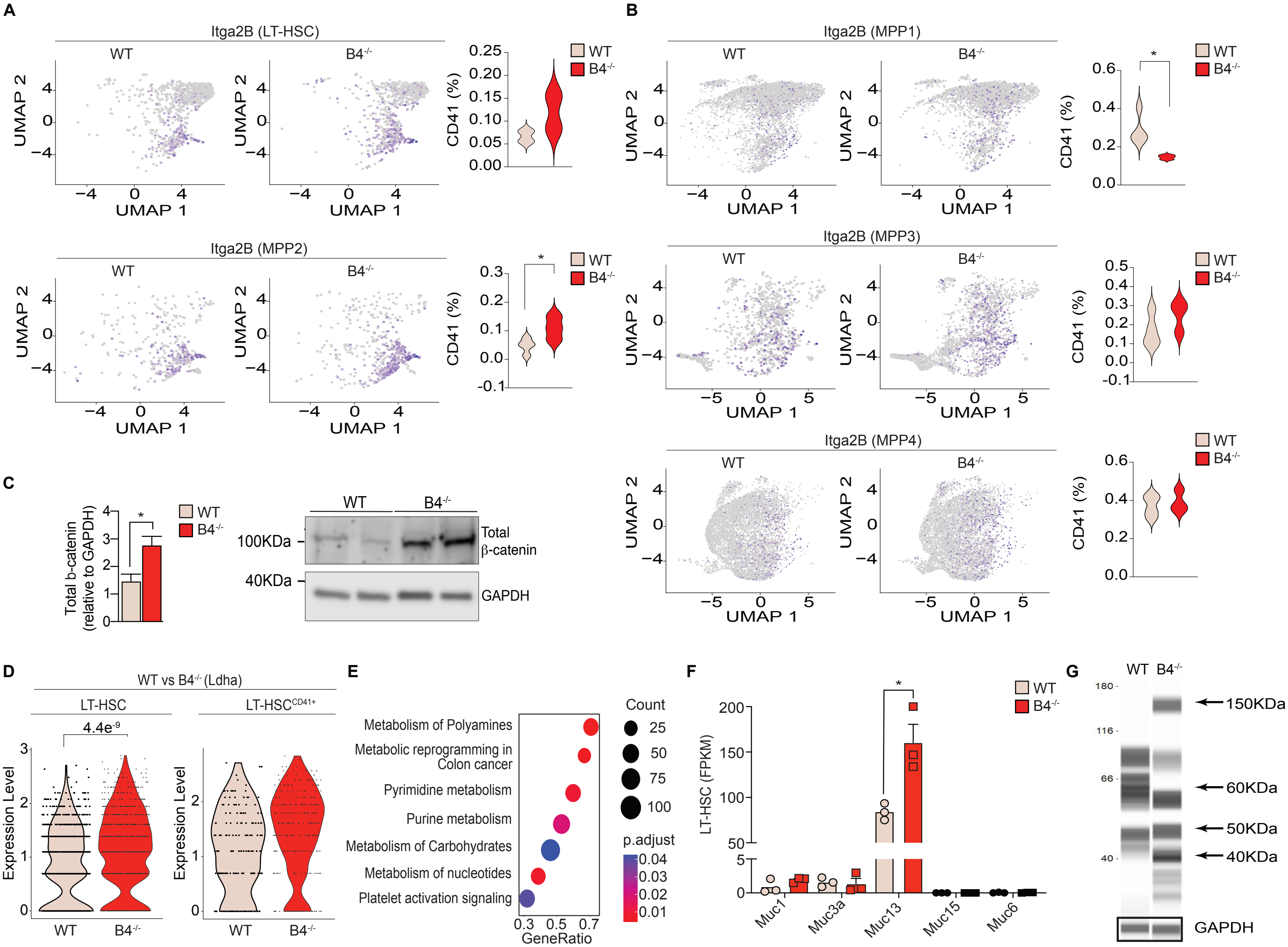

### Figure S5

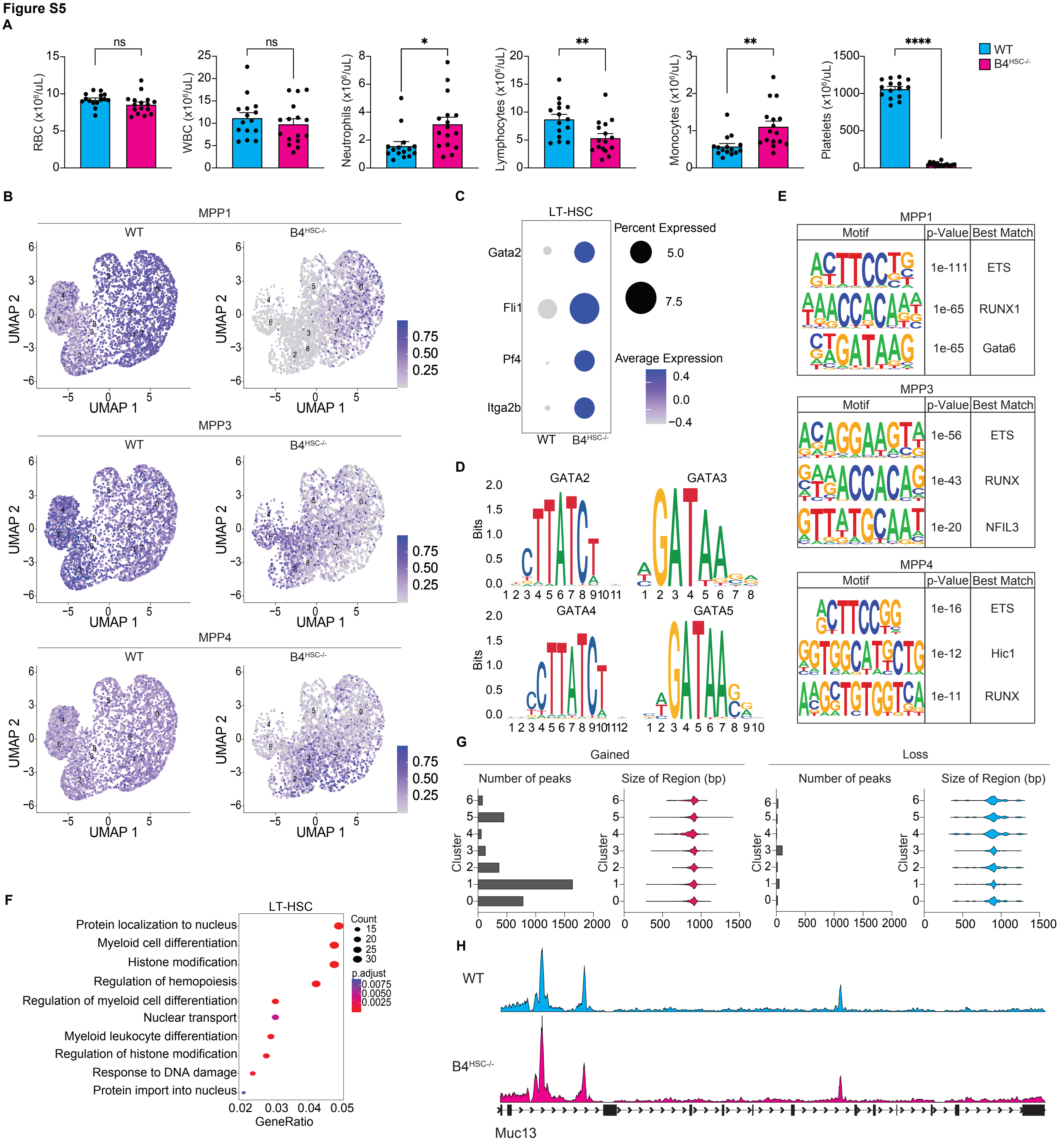
